# Positive and Neutral Updating Reconsolidate Aversive Episodic Memories via Different Routes

**DOI:** 10.1101/2021.01.04.424701

**Authors:** Jingyi Wang, Boxuan Chen, Manqi Sha, Yiran Gu, Haitao Wu, Cecilia Forcato, Shaozheng Qin

## Abstract

Aversive memories are long-lasting and prone to have adverse effects on our emotional wellbeing and mental health. Yet, how to remedy the maladaptive effects of aversive memories remains elusive. Using memory reactivation and emotional updating manipulations, we investigated how positive and neutral emotion updates aversive memories for reconsolidation in humans. We found that positive updating after reactivation was equivalent to neutral updating in altering true memories of the aversive story, but introduced more false memory. Moreover, an additional 12 hours of sleep reconsolidation did not further enlarge true memory differences, but attenuated the effect of reactivation and updating on false memory. Interestingly, the neutral rather than the positive updating reduced the emotional arousal of the aversive memory 24 hours later. Our findings provide novel insights into real-world therapeutic applications regarding how updating with positive and neutral emotion may reshape aversive memories, especially when taking wake- and sleep-filled reconsolidation into account.

## Introduction

Emotional episodic memory constitutes our individual lives. While joyful moments sparkle our memory bank with happiness, there are also incidents of aversive and distressing events, even traumatic. Our memories for the latter once become maladaptive, could burden our emotional wellbeing and mental health. For example, the sudden outburst of the global COVID-19 pandemic has infected over 80 million people and led to more than one million deaths. The part-for-ever caused by this pandemic has become traumatic memories of millions of people worldwide (Rajkumar, 2020). Scientists have warned that overloaded traumatic episodic memory could lead to severe mental disorders like post-traumatic stress disorder (PTSD) and major depression even in post-pandemic time (Carmassi et al., 2020; Kathirvel, 2020; Xiao, Luo, & Xiao, 2020). Appropriate and efficient interventions for maladaptive memory to prevent the development of severe mental disease are in need. There are pieces of evidence showing pharmacological manipulations like introducing adrenergic receptor antagonist (Lonergan, Brunet, Olivera-Figueroa, & Pitman, 2013; Soeter & Kindt, 2015), and electroconvulsive therapy (ECT) might interference with aversive episodic memory (Kroes et al., 2014; Misanin, Miller, & Lewis, 1968). However, these methods are limited for applications due to their aggressive nature. Hence mild and patient-friendly approaches, like behavioral modifications, are under the urgent need for real-world applications.

It has been recognized that episodic memory is amenable. Convincing evidence has proved that after memory retrieval, already consolidated memory can be rendered to a labile state; thus a *reconsolidation* process is required to stabilize it into the brain again (Nader & Hardt, 2009). Once a memory is destabilized, it becomes susceptible to new information, leading to the existence of a time window for memory updating (Phelps & Hofmann, 2019). The discovery of memory malleability has generated broad interest to remedy symptoms linking to aversive memories in clinical populations (i.e., PTSD and depressive patients). Emerging evidence has suggested that introducing new learning shortly after memory reactivation can incorporate new information into the already labialized original memory, a process known as memory updating (Lee, Nader, & Schiller, 2017). Among conventional memory updating approaches, counter-conditioning that involves replacing an expected salient outcome with a new outcome of the opposite valence (Keller, Hennings, & Dunsmoor, 2020) has been proved with promising outcomes for appetitive-to-aversive applications. For instance, introducing aversive feelings shortly after memory reactivation had significantly reduced addictive behavior for food (Olshavsky et al., 2013), alcohol (Das, Lawn, & Kamboj, 2015), and cocaine (Goltseker, Bolotin, & Barak, 2017). On the contrary, aversive-to-appetitive counter-conditioning studies reported inconsistent findings. On the one hand, utilizing behavioral and optogenetic techniques, researchers had successfully remedied depressive-like or aversive behavior in rodents, by artificially triggering positive memory engrams in the hippocampus during negative experiences (Ramirez et al., 2015; Redondo et al., 2014). On the other hand, findings from the aversive-to-appetitive counter-conditioning paradigm in humans were inconsistent. While many studies have shown a better effect for aversive-to-positive updating than aversive-to-neutral updating or extinction (Eifert, Craill, Carey, & O’Connor, 1988; Newall, Watson, Grant, & Richardson, 2017; Reynolds, Field, & Askew, 2018), there are also studies showing that counter-conditioning is not better than the traditional extinction protocol in some aspects (de Jong, Vorage, & van den Hout, 2000; Meulders, Karsdorp, Claes, & Vlaeyen, 2015), and the counter-conditioning could even prone to renewal the negative memory (Holmes, Leung, & Westbrook, 2016). Hence, although with promising real-world application potentials, the experimental boundary of counter-conditioning updating remains to be explored.

Episodic memory for an emotional event consists of multiple aspects of memory contents, including episodic details on what, where, and when such event occurred (Tulving, 1993), as well as the emotional significance of how we feel about that experience (Christianson, 1992; Dolan, Lane, Chua, & Fletcher, 2000). The recall of episodic memory includes the information that one truly experienced (i.e., true memory), and some fictitious yet plausible information that did not experience (i.e., distorted or false memory) (Guarnieri, Bueno, & Tudesco, 2019; Loftus, 1979). The fuzzy-trace theory (FTT) proposes that the specific and detailed traits of the episodes are stored in a literal memory system, the recalling of which would generate true memory. In contrast, the gist and central theme of the episodes are stored in the essence memory system, generating false memory (Brainerd & Reyna, 2002, 2005). Besides the memory contents, the subjective emotional feeling is a vital component of emotional episodic memory, and it is also a crucial indicator to evaluate therapeutic modifications (Lane, Ryan, Nadel, & Greenberg, 2015). Ample studies have successfully reduced the individual’s negative feelings (e.g., fear) through memory updating operations (Sandkühler & Lee, 2013; Soeter & Kindt, 2015). However, to our knowledge, there is no study yet to investigate how updating with positive emotion reshapes previously acquired aversive memories with either impaired or distorted outcomes, and even less is known how this procedure alters the emotion of original memory representation.

Memory reconsolidation involves protein synthesis processes, requiring a period to complete (Nader, Schafe, & LeDoux, 2000). Some studies suggested that the reconsolidation window lasts at least six hours during wakefulness (Björkstrand et al., 2016). Meanwhile, it is also reported that sleep supports memory reconsolidation (Klinzing, Rasch, Born, & Diekelmann, 2016; Walker, Brakefield, Hobson, & Stickgold, 2003), probably by speeding up or shortening the reconsolidation window (Moyano, Diekelmann, Pedreira, & Forcato, 2019). This indicates that both sleep and wake play an essential role in memory reconsolidation. In the context of emotional episodic memory, it is well known that sleep is crucial to consolidate newly encoded episodic events for both true memories (Weber, Wang, Born, & Inostroza, 2014) and false memories (Payne et al., 2009). Importantly, when combining the modulation of emotion, sleep seems to consolidate these two kinds of memory in different ways (McKeon, Pace-Schott, & Spencer, 2012). Reconsolidation works as a follow-up step after memory retrieval rather than encoding, and it naturally involves both wake and sleep. Hence it is interesting to ask how sleep, after the wake reconsolidation is completed, continually supports reconsolidation for both the true and false memory of episodic events that have been emotionally modified. To our knowledge, this has not been well studied.

In the present study, we attempted to close the research gaps mentioned above by asking the following questions: First, whether new information introduced shortly after reactivation can alter existing aversive memory under the reconsolidation procedure? Second, whether introducing interference with positive emotion can impair previous negative memory better than updating with neutral emotion? And third, how the effects of positive and neutral updating on aversive memory evolve over 12-hour wakefulness and 24-hour interval with a night of sleep? We conducted a between-subject factorial experiment to address these critical questions, including the independent variables of reactivation and positive/neutral updating procedures. Participants encoded an aversive story presented as picture slides with auditory narratives on the Day1 morning, with an immediate recall test as the baseline assessment of memory (**Figure 1**). Seven days later (on the morning of Day 8), participants returned to the lab in morning and were assigned into four experimental groups with either memory reactivation combining with positive or neutral updating or just updating manipulations (Group A, ReaPos; B, ReaNeu; C, NReaPos; D, NReaNeu; see *Methods* for detailed descriptions). For the method of reactivation, it is known that prediction errors (or surprise) lead to destabilization of memory traces (Exton-McGuinness, Lee, & Reichelt, 2015; Sinclair & Barense, 2018). Hence we falsely claimed a memory retrieval session to the participants but surprisingly interrupted the retrieval after the first cue slide was presented (see *Methods*), the way that had been used for successful memory destabilization (Forcato et al., 2007). All participants were tested 12 hours later in the evening of the same day for their memory of the original aversive memory. Finally, after a night of sleep (on the morning of Day 9), all participants returned to the lab in the morning for a final test on memory, emotional valence, and arousal of the negative story.

**Figure 1:**
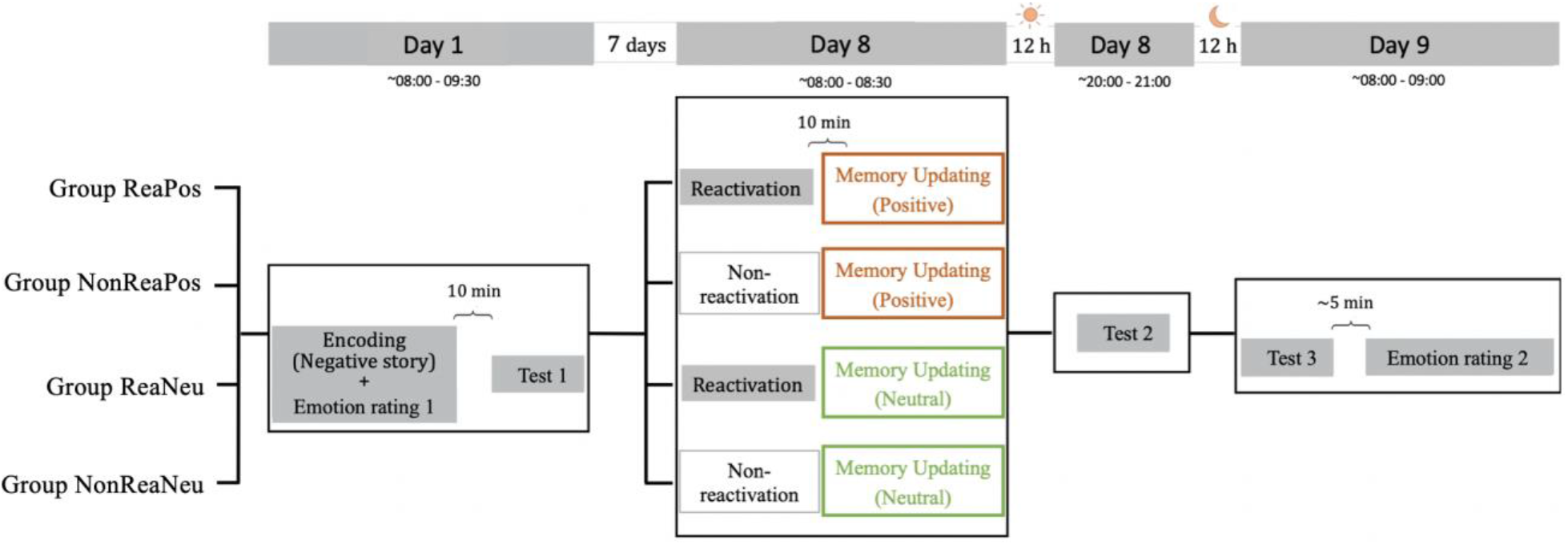
Experimental design. Participants were assigned to one of four groups (A, B, C, D). On the morning of Day 1, participants from all groups were shown a negative story presented in slides. Participants were asked to rate their emotional valence and arousal for the presented slide and the whole story (Emotion rating 1). Ten minutes after the encoding, the participants orally recalled the content of the just-viewed story (Test 1). On the morning of Day 8, participants from Group A and C were told to recall the negative story but were interrupted by the experimenter on purpose to produce memory reactivation. Participants of Group B and D did not experience the memory reactivation process. Ten minutes later, participants in groups A and B received positive memory updating, while groups C and D received a neutral updating. Twelve hours later in the evening, all the participants recalled the negative memory again (Test 2). After a night of sleep, all of them recalled the negative memory on the morning of Day 9 (Test 3). Five minutes later, all participants scored their emotional valence and arousal of the original negative story again (Emotion rating 2).

With this design, we hypothesized that 1) an impairment of memory for the original negative story on Day-8 only appears in the groups that received both memory reactivation and updating; 2) Since the strength of the updating material differs the degree of impairment for the old memory, i.e., stronger new learning after reactivation leads to better memory updating, we would expect that the group with positive emotion updating should impair the negative memory more than the neutral group. Moreover, 3) a recent study in starlings showed that sleep promoted the reconsolidation of old memory once it has been reactivated, and this effect was notably more substantial when new interferences existed after the reactivation (Brawn, Nusbaum, & Margoliash, 2018). Hence, we hypothesize that after 24-hour interval involving sleep could favor the recovery of the negative memory, especially in the positive group (as a stronger updating manipulation than the neutral updating) with memory reactivation. Apart from memory impairment, an early study suggested that after reactivation, the destabilized memory turns to be more susceptible to interference, intermixing with new information (Hupbach, Gomez, Hardt, & Nadel, 2007). Thus, we further hypothesized that 4) comparing with non-reactivation groups, memory performance from reactivation groups should have more false memory after the updating stories. Finally, we hypothesized that the emotional components of the updating material should also be integrated into the negative memory after reactivation, leading to a significant reduction of the negative feeling of the aversive memory for subjects who received a positive updating.

## Results

### Memory updating effect of the negative story after 12-hour wakefulness

First, we investigated the effects of reactivation and the emotional type of updating on episodic memory, including true and false memory aspects of the initial negative story. Ten minutes after initial encoding, there were no significant differences for neither true memory (all *F*_(1, 76)_ < 2.385, all *p* > 0.127), nor false memory performance (all *F*_(1, 76)_ < 2.428, all *p* > 0.123) across four experimental groups. Because the self-paced unrestricted free recall naturally generated large variance between participants (True Memory score range: 27-232.5; False Memory score range: 4-135), between-group difference could be masked by the high variance within each group. Therefore, we normalized the memory performance of the two delayed recalls by dividing their performance on the previous test (Day8/Day1) of each group to measure memory changes. We conducted two separate 2 (Reactivation: Reactivated vs. non-Reactivated) × 2 (Updating: Positive vs. Neutral) between-subjects ANOVAs for true and false memory scores. For true memory, this analysis revealed a strong main effect on Reactivation (*F*_(1, 76)_ = 13.773, *p* < 0.001, η^2^ = 0.153), which means non-reactivated groups remained similar recall level as their performance on Day 1, in contrast, the reactivated groups of each updating type remembered less true information of the negative story on Day 8 than Day1 (**Figure 2A**). However no significant interaction effect for neither Reactivation × Updating nor main effect of Updating (all *F*_(1, 76)_ < 0.182, all *p* > 0.613). This indicated that the true memory performance of the negative story in the reactivated groups is significantly worse than the non-reactivated groups, independent of whether they were updated with positive or neutral emotion. On the contrary, the percentage of false memory changes on Day 8 relative to Day1 showed a trending significant interaction effect (*F*_(1, 76)_ = 3.724, *p* = 0.057, η^2^ = 0.047) (**Figure 2B**).

**Figure 2.**
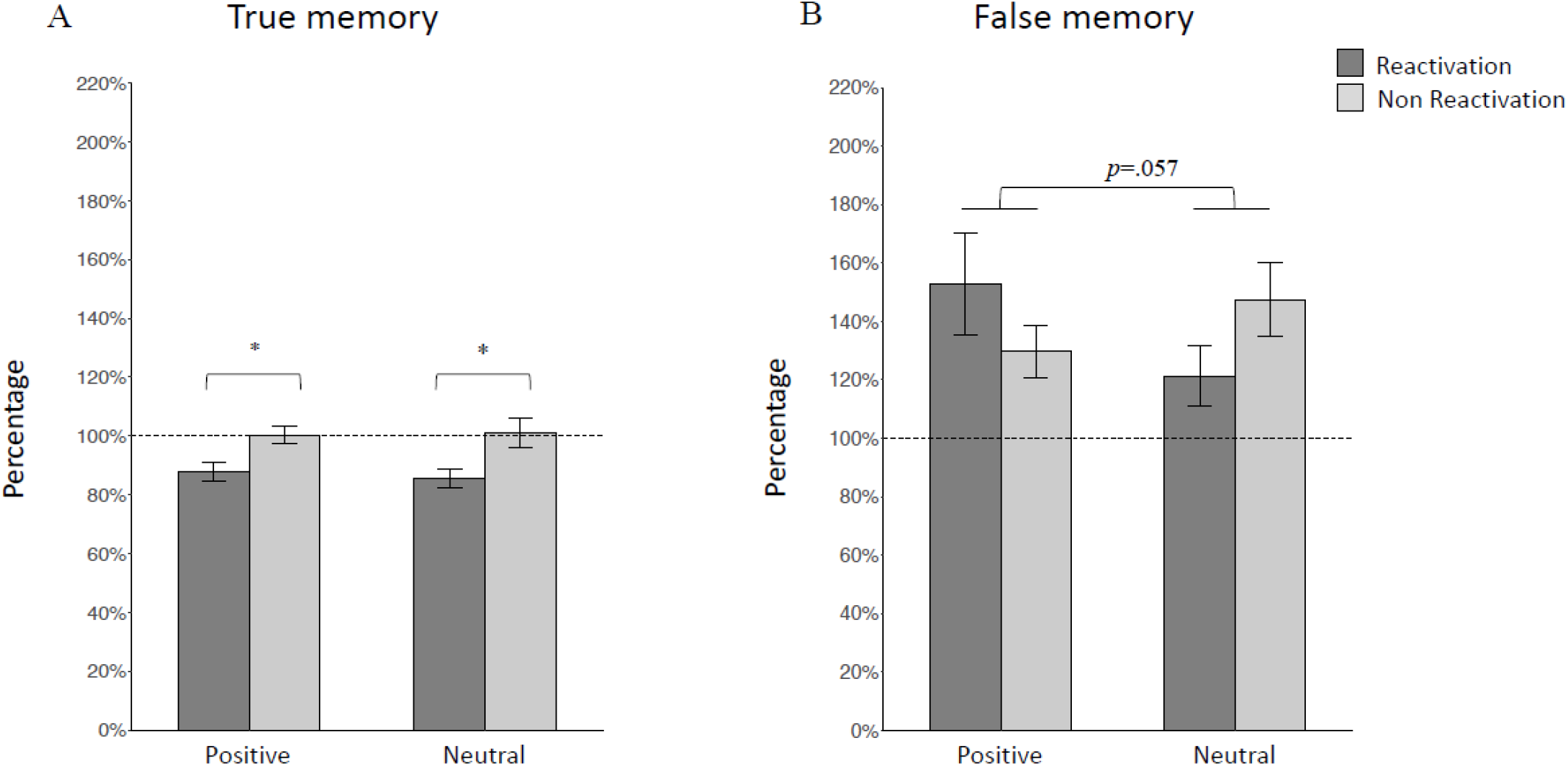
Memory performance for true and false memory at Test 2 on Day 8. Performance is indicated as the percentage of recalled correct (A) or incorrect (B) information on Day 8 with their corresponding performance on Day 1 set to 100%. Asterisks represent p-values from post hoc simple effect analysis and interaction effect of ANOVA in Figure 2A and 2B, respectively. Percentage for correct (A) information per group: Rea&Pos group: 88.0% ± 3.2%. NonRea&Pos group: 100.2% ± 3.0%. Rea&Neu group: 85.7% ± 3.2%. NonRea&Neu group: 101.1% ± 5.2%. Percentage for incorrect (B) information per group: Rea&Pos group: 152.7% ± 78.2%. NonRea&Pos group: 129.6% ± 39.7%. Rea&Neu group: 121.3% ± 45.9%. NonRea&Neu group: 147.4% ± 56.4 %. Data are mean ± S.E.M. Rea: reactivation. Pos: positive. Neu: Neutral *: p < 0.05.

### Memory updating effect of the negative story after 24-hour interval including sleep

Next, we further investigated the updating effects on the negative memory after a 24-hour interval, as a typical reconsolidation window. Again, we conducted a 2 (Reactivation: Reactivated vs. non-Reactivated) × 2 (Updating: Positive vs. Neutral) between-subjects ANOVAs on memory performance of Day 9 relative to Day 1. It revealed a significant main effect of Reactivation for true memory (F_(1,76)_ = 7.19, *p* = 0.009, η^2^ = 0.086). Similar to the effect on Day 8, we observed significantly less true memory on Day 9 relative to Day 1 for the reactivation than the non-activation groups regardless of positive and neutral updating conditions(**Figure 3A**). But, a parallel 2-by-2 ANOVA revealed a non-significant interaction effect (F_(1, 76)_ = 3.024, *p* = 0.086, η^2^ = 0.037) of false memory (**Figure 3B**).

**Figure 3.**
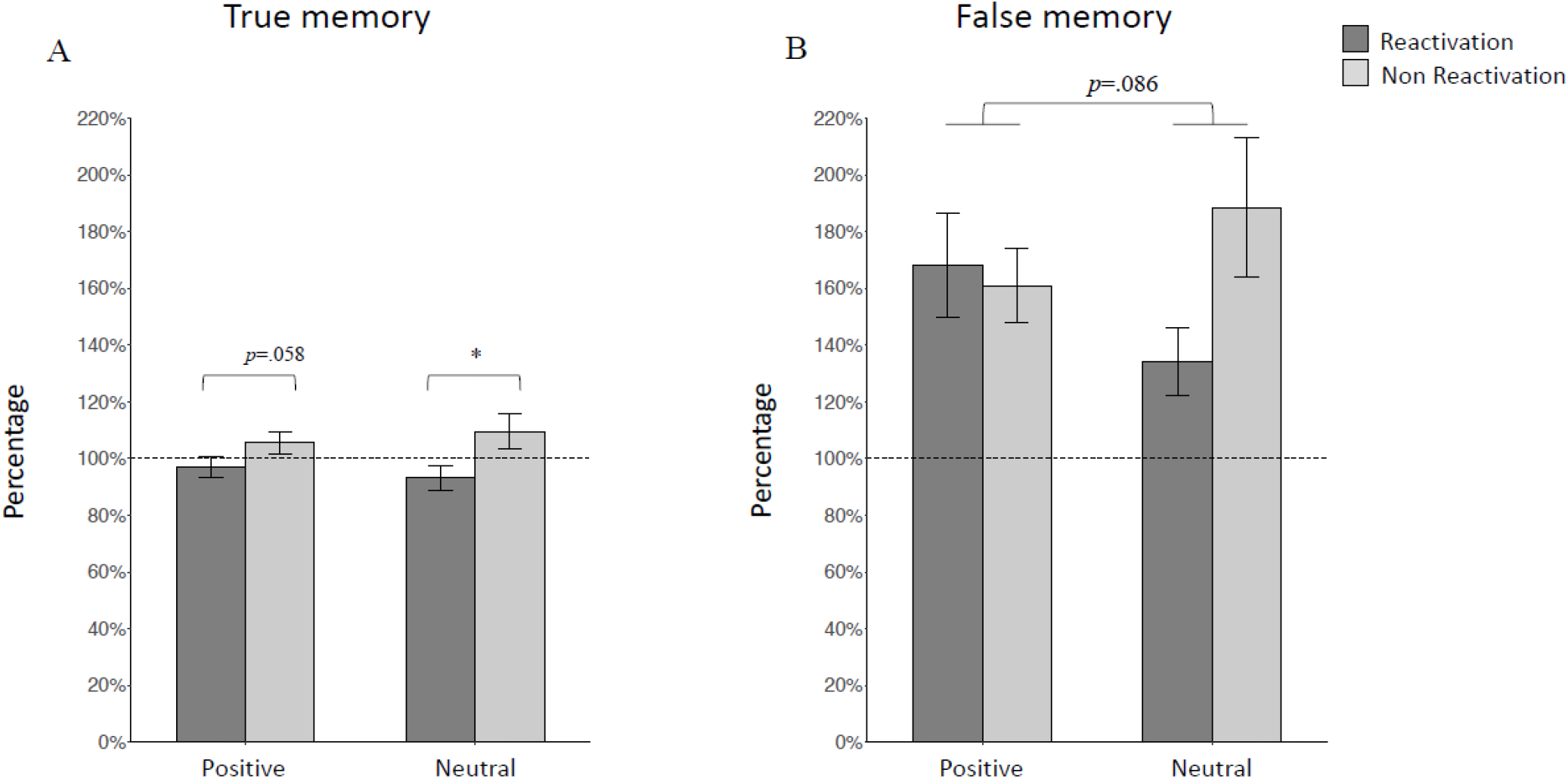
Memory performance for true and false memory at Test 3 on Day 9. Performance is indicated as percentage of recalled correct (A) or incorrect (B) information on Day 9 with their corresponding performance on Day 1 set to 100%. Percentage for correct (A) information per group: Rea&Pos group: 97.0% ± 15.6%. NonRea&Pos group: 105.5% ± 18.0%. Rea&Neu group: 93.2% ± 19.4%. NonRea&Neu group: 109.6% ± 27.8%. Percentage for incorrect (B) information per group: Rea&Pos group: 168.3% ± 82.2%. NonRea&Pos group: 161.0% ± 59.0%. Rea&Neu group: 134.1% ± 53.2%. NonRea&Neu group: 188.4% ± 110.1%. Data are mean ± S.E.M. *: p < 0.05.

For further investigation about the different changes between the true memory and false memory going through a night of sleep, we conducted a 2 (Reactivation: Reactivated vs. non-Reactivated, between factor) × 2 (Updating: Positive vs. Neutral, between factor) × 2 (Sleep: pre-sleep vs. post-sleep, within factor) repeated ANOVA on true and false memory performance of Day 8 and Day 9 relative to the memory on Day 1(i.e., Day 8/Day 1 and Day 9/Day 1). After a night of sleep, we observed a memory increase (all *F* > 35.122, all p <0.001) for both true and false memory. While there was no any significant interaction effect for the change of true memory across sleep (all *F*_(1,76)_< 1.357, all *p* > 0.235), the change of false memory showed a significant interaction effect between Sleep and Reactivation (*F*_(1, 76)_ = 6.662, *p* = 0.012, η^2^ = 0.081, **Figure 4**). Post-hoc tests for simple effect revealed that this significant difference was mainly driven by the difference between before and after sleep within the non-reactivated groups (t_(1, 76)_ = 6.016, *p* < 0.001). In order to reveal the interaction effect between sleep and reactivation of the false memory performance on Day 9 relative to Day 8, the present study built a further 2 (Reactivation: Reactivated vs. non-Reactivated, between factor) × 2 (Updating: Positive vs. Neutral, between factor) ANOVA revealed a main effect on Reactivation (*F*_(1, 76)_ = 3.822, *p* = 0.044, η^2^ = 0.052, **Figure 5**), with the reactivated groups increased less false memory than the non-reactivated groups. These results indicate that true memory of the negative story rises evenly and slightly in all four groups after a night of sleep. However, memory reactivation prohibited false memory to increase more over 24-hour interval involving overnight sleep. This change after one-night of sleep blurred the interaction effect between reactivation and updating of false memory.

**Figure 4.**
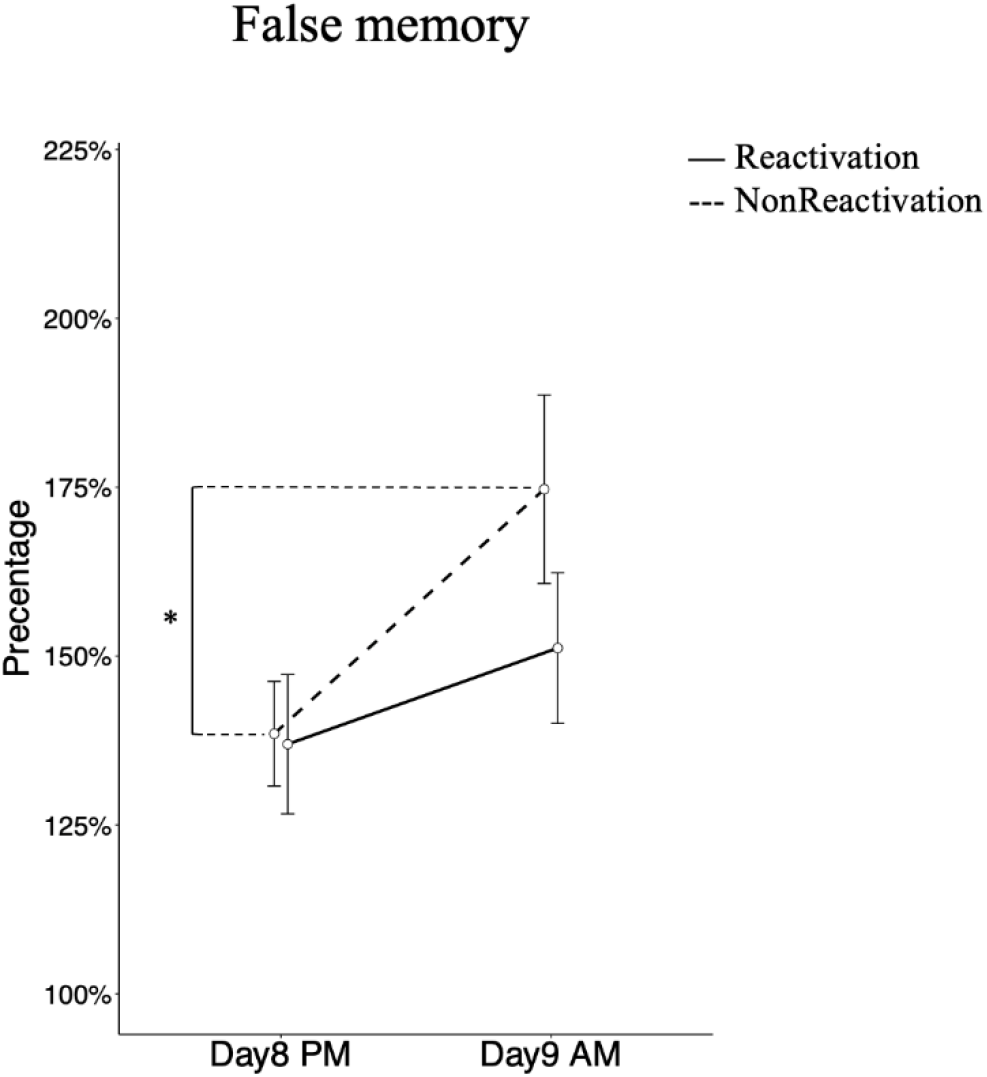
Memory performance for false memory at Test 2 on Day 8 and Test 3 on Day 9. Performance is indicated as the percentage of incorrect information before and after sleep with their corresponding performance on Day 1 set to 100%. Percentage for incorrect information per group: Rea&Day8: 137.0% ± 11.1%. NonRea&Day8: 138.5% ± 14.0%. Rea&Day9: 151.2% ± 11.1%. NonRea&Day9: 174.7% ± 13.4%. Percentage of neutral updating groups. Data are mean ± SEM.

**Figure 5.**
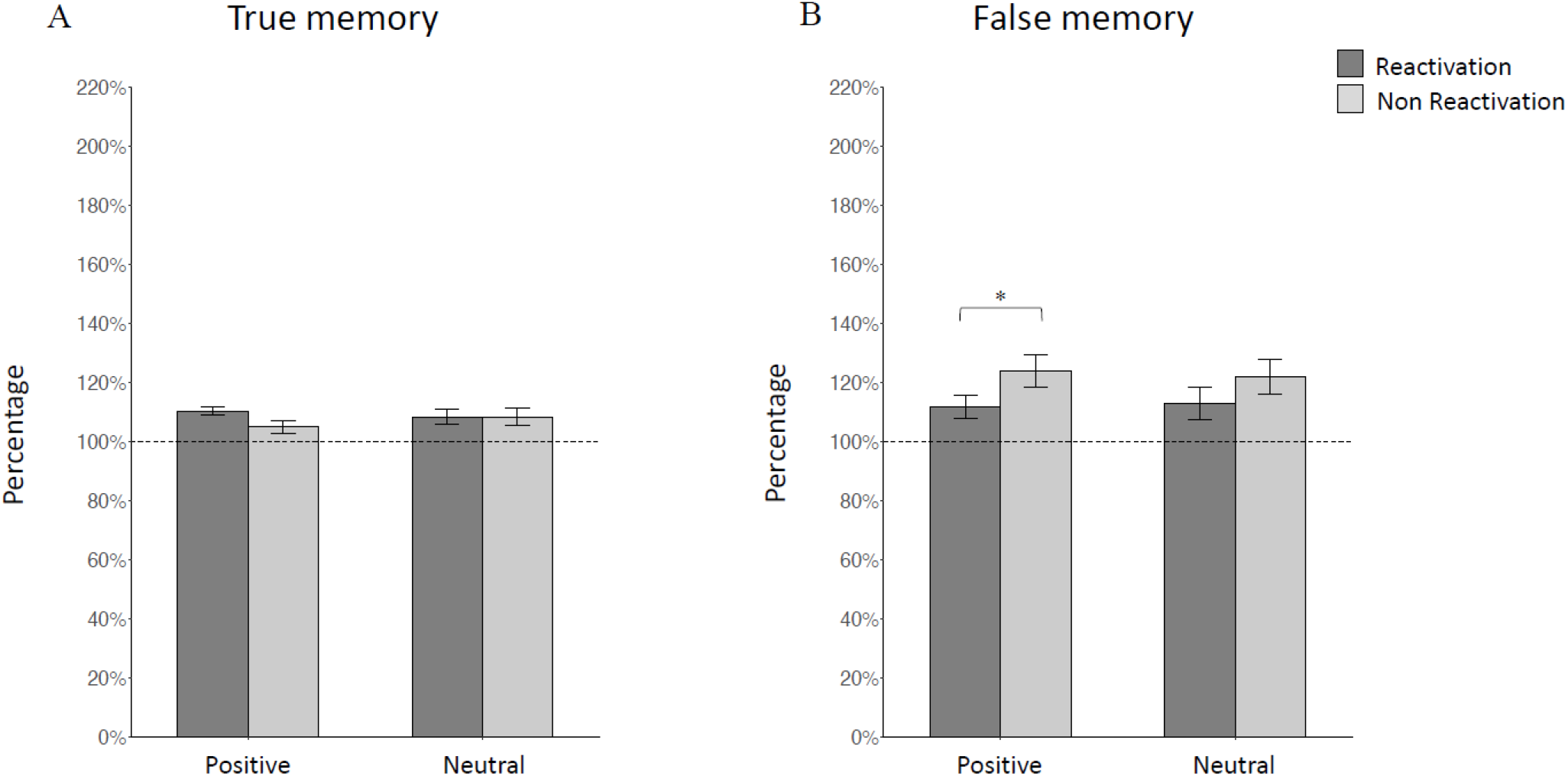
Memory performance for true and false memory at Test 3 on Day 9. Performance is indicated as percentage of recalled correct (A) or incorrect (B) information on Day 9 with their corresponding performance on Day 8 set to 100%. Percentage for correct (A) information per group: Rea&Pos group: 110.4% ± 5.7%. NonRea&Pos group: 105.0% ± 9.2%. Rea&Neu group: 108.4% ± 11.9%. NonRea&Neu group: 108.5% ± 12.4%. Percentage for incorrect (B) information per group: Rea&Pos group: 111.7% ± 17.2%. NonRea&Pos group: 124.0% ± 24.3%. Rea&Neu group: 112.8% ± 24.6%. NonRea&Neu group: 121.9% ± 26.2%. Data are mean ± S.E.M. *: p < 0.05.

### Neutral, rather than positive updating, altered emotional arousal of the negative story after reconsolidation

Besides the changes in true and false memory, we also investigated whether introducing an opposite emotion after memory reactivation can alter the original aversive memory’s emotional feelings. Unlike memory that is based on original encoded information with multiple tests on Day 8 and Day 9, the feeling of an event is caused mainly by the perception of the current events(Schachter & Singer, 1962). Hence we only obtained the emotional valence and arousal subjecting to the ratings on Day 9, rather than the changes over time. A 2-by-2 ANOVA with Reactivation and Updating as between-subject factors revealed that the subjective rating of valence for the negative story was not different between experimental groups, neither on Day 1 nor on Day 9 (all *F*_(1, 76)_ < 1.548, all *p* > 0.217). However, the ratings of arousal on Day 9 showed a significant interaction effect (*F*_(1, 76)_ = 8.12, *p* = 0.006, η^2^ = 0.096), with higher scores for the positive reactivated than positive non-reactivated groups and neutral reactivated groups (**Figure 6**). To note, although we did not observe any significant difference for the arousal on Day1 (all *F*_(1, 76)_ < 2.463, all *p* > 0.143), it was strongly correlated with the arousal on Day 9 (r = 0.644, *p* < 0.001, Spearman’s rank correlation coefficient), suggesting some consistencies of the rating across the whole experiment. Hence, we introduced the arousal rating on Day 1 as a covariate variable, and the interaction we previously observed on Day 9 still held true (interaction effect: *F*_(1, 76)_ = 6.234, *p* = 0.015, η^2^ = 0.039). Post hoc analysis for simple main effect on the interaction effect revealed that this significant difference was mainly driven by the difference between the reactivated groups with positive vs. neutral updating (t_(1, 76)_ = 2.617, *p* = 0.011), and the difference between the two positive updating groups (t_(1, 76)_ = −2.33, *p* = 0.023).

**Figure 6.**
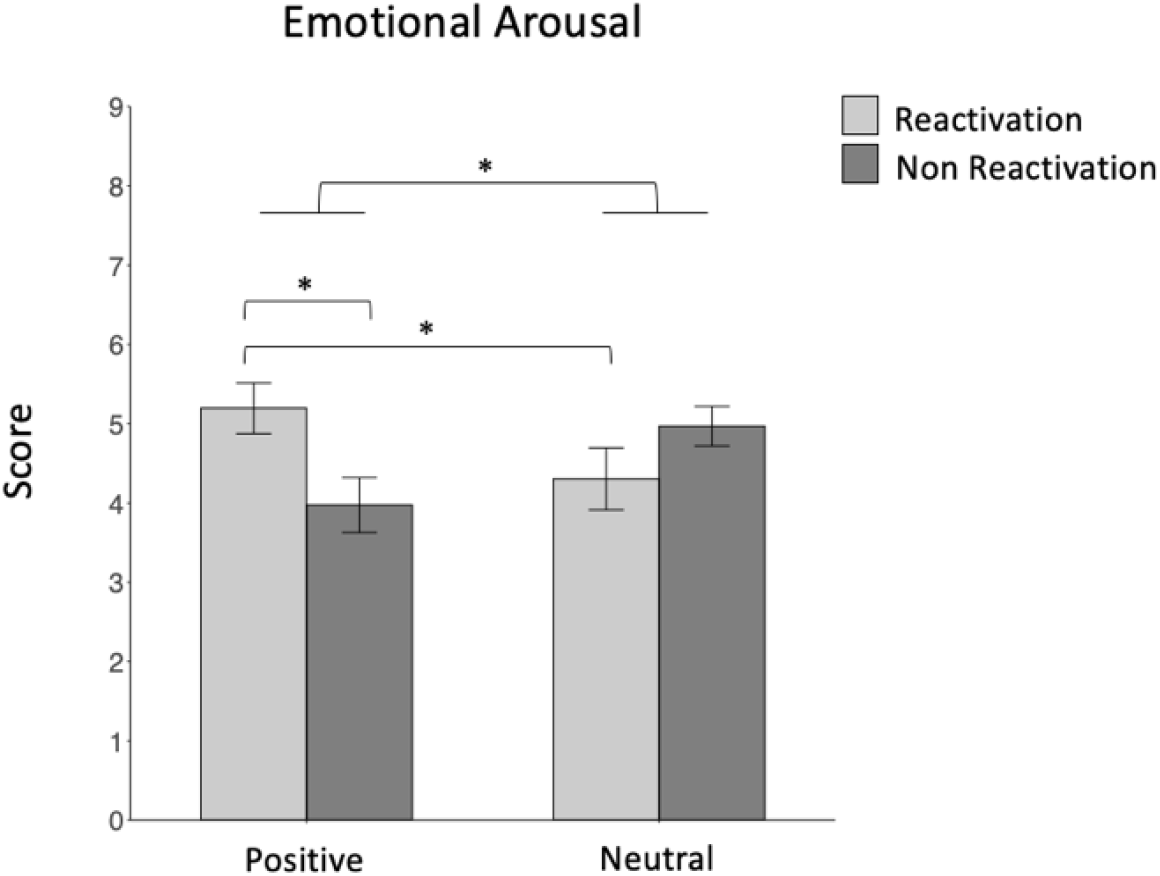
Subjective ratings of emotional arousal on Day 9. Y-axis is the emotional arousal score on Day 9 (larger values indicated higher arousal). Score per group: Rea&Pos group: 5.195 ± 1.428. NonRea&Pos group: 4.132 ± 1.426. Rea&Neu group: 4.589 ± 1.599. NonRea&Neu group: 4.928 ± 1.126. Data are mean ± S.E.M. *: p < 0.05.

It is worth to note that none of the control measures describing participants’ general emotional state (subjectively rated anxiety, depression, and general stress level) showed any significant differences across the whole experimental procedure for the four experimental groups (all *F*_(1, 76)_ < 1.706, all *p* > 0.174). Meanwhile, participants were also comparable among groups in empathy (*F*_(1, 76)_ = 1.003, *p* = 0.397), and the PVT scores (all *F*_(1, 76)_ < 0.523, all *p* > 0.200), suggesting the changes of the episodic memory is unlikely due to the differences from the sense of substitution and tiredness of each participant.

## Discussion

In the present study, we investigated how positive, and neutral updating after memory reactivation reshapes previously acquired aversive episodic memory after reconsolidation over 12-hour wakeful and 24-hour (including sleep) intervals. As expected, memories from the reactivated groups were reduced on Day 8 compared with non-reactivation groups, suggesting that memory reactivation rendered original memory susceptible to new information. The updating effect was detectable after a 12-hour wake window followed reactivation. However, following memory reactivation, introducing positive emotion did not impair the memory of the learned initially negative story more than the ones updated with neutral information but induced more false memory, similar as it was observed in the neutral updating without reactivation. Moreover, one-night sleep did not seem to alter the reconsolidation outcomes for true memory within the 12-hour wake period followed reactivation, whereas it blurred the reconsolidation effect on false memory. Finally, the neutral updating rather than the positive updating after memory reactivation, surprisingly, reduced emotional arousal of the negative story 24 hours later. In contrast, the valence of the negative story was comparable among groups.

### Emotional updating for aversive memories after 12-hour wake intervals

Episodic memory for the negative story was impaired only for the groups with memory reactivation (relative to non-reactivation) on Day 8, regardless of positive and neutral updating, indicating that our reactivation manipulation (i.e., arbitrarily interrupting the retrieval process to elicit prediction error) was successful in destabilizing the original memory trace (Kindt, Soeter, & Vervliet, 2009). This pattern of results is in line with previous studies on declarative memory reconsolidation in humans (Forcato, Argibay, Pedreira, & Maldonado, 2009; Forcato et al., 2007; Hupbach et al., 2007). Memory reactivation was believed as an efficient way to render memories malleable. In the context of memory reconsolidation, memory reactivation is usually effectively induced by prediction errors, a mismatch of the expected outcome with a surprising outcome. It has been suggested that phasic activity of midbrain dopamine neurons (particularly in the ventral tegmental area, VTA) is considered to represent the prediction error that drives learning (Reichelt, Exton-McGuinness, & Lee, 2013; Sinclair & Barense, 2019), subsequently trigger destabilization of existing memory traces, and thus drives new learning (Exton-McGuinness et al., 2015). Our observed reactivation effect on true memory is consistent with a recent study that has also extended the prediction error engagement in episodic memory reconsolidation. By violating the action-outcome of video clips to generate the feeling of surprise (i.e., prediction error), this study showed that the memories that have been surprisingly interrupted were more susceptible to subsequent interference from a new set of semantically related videos, and the more surprising the subject rated, the more intrusion were induced. Our present study supports this research, showing that memory updating, most likely via introducing a prediction error, is a prerequisite to interrupt previously acquired aversive episodic memory.

After memory reactivation, updating with positive emotion did not alter true memories of the negative story when compared with the neutral updating. Instead, it yielded more false memories after 12 hours. In other words, integrating positive emotion into a destabilized aversive memory was similar to integrating with neutral emotion in impairing the recall of episodic details but generated more fictitious episodic details than neutral updating. Human memory is never a loyal record of the past, and the generation of false memory has also been view as an adaptive way to enable memory flexibility to adapt to new situations for ever-changing cognitive and environmental needs (Sinclair & Barense, 2018). Why insert a positive emotion right after memory destabilization did not further impair true memory but accelerated memory distortions in later memory testing? A potential explanation could refer to the *affect-as-information theory* by Schwarz Norbert, which suggests that an individual’s affective state may influence their style of information processing (Schwarz, 1990). According to this theory, when an individual is in a bad mood, their reasoning tends to be “low degree of originality, creativity, and playfulness”; thus, they are likely to recall the literal (i.e., verbatim) memories. However, when in a positive state, one tends to believe his or her current environment “is safe, hence tends to use heuristics in information processing,” thereby they are more likely to use gist-based memory recall strategies. This tendency may reduce memory load (for example, store fewer details) and support retention of themes and meanings that facilitate generalization and abstraction. However, it could also open opportunities to generate false information due to the involvement of constructive processing (Schacter, Guerin, & Jacques, 2011). In other words, individuals with positive emotion might prefer gist-based processing to reconstruct the reactivated memories, thus producing more gist-based generalization based on their memories. Indeed, gist-based generalization has been linked to false memories for related events by many behavioral studies in humans (Guerin, Robbins, Gilmore, & Schacter, 2012; Koutstaal & Schacter, 1997; Parker & Dagnall, 2007).

In the present study, the valence of updating stories was rated by participants significantly different between neutral and positive, indicating that the participants well distinguished the two emotional states after viewing the updating stories. Therefore, during the updating session, either positive or neutral emotion was then integrated into the already unstabilized negative memory (due to memory reactivation). According to the *affect-as-information* theory, we thus suspect that participants in positive emotion (i.e., encoding a positive story) could believe that the environment is safe, and hence selectively promoted heuristic and gist-based (i.e., relational) processing, which resulted in more memory errors (Schwarz, 1990). On the other hand, true memory was the memory of the real retrieval of detailed information that happened in an experienced event, which, according to Schwarz Norbert, was more sensitive to negative rather than positive mood (Schwarz, 1990). Although we only set up positive versus neutral (rather than negative) updating in our current design, a similar mechanism might still be relevant to explain why we did not observe any true memory difference caused by the type of emotion for updating. Future studies with more targeted designs are needed to address this speculation.

### Emotional updating on aversive memories after 24-hour intervals, including sleep

Reconsolidation is a long-lasting process, including wake and sleep (Dudai, 2012). Our present study speculated that the longer delay after reactivation and updating, including sleep, could transform true and false memory in different mechanisms. Indeed, we observed differential changes in true and false memory over the 24-hour interval from Day 8 to Day 9. For true memory, as discussed above, the reactivation combined with updating disturbed the original negative memories already after a 12-hour wake delay (i.e., in the Day 8 evening). And this effect did not further evolve during sleep as the four groups did not show any significant differences over sleep (i.e., Day 9 morning). Several previous studies have suggested that sleep typically facilitates memory reconsolidation (Klinzing et al., 2016; Moyano et al., 2019), and some changes from reconsolidation can only be evident after a period filled with sleep (Kindt & Soeter, 2018; Simon, Gómez, & Nadel, 2020). However, reconsolidation is not the same procedure as the original consolidation, but having a shorter time window (Debiec, LeDoux, & Nader, 2002; Phelps & Hofmann, 2019). Hence, a period of wakefulness could be already sufficient to complete the reconsolidation procedure (Björkstrand et al., 2016). Thus the following sleep could function to stabilize the changes accomplished during wakefulness. Paying attention to wake and sleep delay progressively, Our results refined the reconsolidation process into steps and inspired that 12 hours of wakefulness per se was sufficient to reconstruct true memory from reactivation and updating. However, lacking direct comparison with immediate sleep after the updating, we cannot conclude whether true memory reconsolidation is really wake-rather than sleep-dependent. Nevertheless, due to the evidence that sleep (or even a short nap) seems to shorten the reconsolidation window, by showing comparable effects as with a long wake reconsolidation (Moyano et al., 2019), we would speculate that sleep right after the memory updating could show similar results as we reported here with the wake-sleep interval. Further sleep reconsolidation could still be necessary to retain the ultimateness of reconstruction, thus endure the reconstructed memory continually. This speculation cannot be proved from the present study due to the lack of direct comparison with a wake control group. However, this possible mechanism could enlighten us to have a more in-depth understanding of the observed time-dependent memory reconsolidation dynamics.

Unlike true memory, we observed the impact of sleep on false memory that was generated during reconsolidation, i.e., false memory in the reactivated groups with both positive and neutral updating increased less oversleep than the non-reactivated groups. This result indicated that sleep promoted memory reconstruction during reconsolidation was more in favor of false memory. It might be because that false memory is relatively more fragile and more reconstructive than true memory with strong bases (Brainerd & Reyna, 2005), which is more likely to profit from sleep (Creery, Oudiette, Antony, & Paller, 2015; Drosopoulos, Schulze, Fischer, & Born, 2007; Oudiette & Paller, 2013) and take longer time to be processed. A study probing into the interaction of sleep and emotional content reported that sleep did not enhance veridical items but fed on false information for emotional materials (McKeon et al., 2012), which was somewhat in line with our results that sleep affects more false rather than true memories. We suspect that it could because the induced positive emotion naturally decayed with time (Davidson, 1998), the intervention of updating in our study might no longer sustain after the 12-hour daytime interval. Hence, both positive and neutral groups embodied their uniform transforming patterns during sleep: the main effect of reactivation in true memory prevailed, and the interaction effect of false memory attenuated. This could be because sleep prefers to preserve the reactivated old memories and thus benefit the reconsolidation of remote memories as previous studies have shown (Klinzing et al., 2016), which could hence maintain true prevent generating more false memory.

In fact, sleep is often relative to more false memory formation with the function of schema integration (i.e., incorporation of initially distinct memories) (Diekelmann, Born, & Wagner, 2010; Diekelmann, Büchel, Born, & Rasch, 2011; Landmann et al., 2014; Payne et al., 2009) and “gist” abstraction and extraction (Durrant & Lewis, 2009; Shaw & Monaghan, 2017). However, empirical studies also support that sleep consolidation reduces false memory by improving recollection of studied details and avoiding false memories, and enabling more efficient and accurate retrieval (Fenn, Gallo, Margoliash, Roediger, & Nusbaum, 2009). Our study is consistent with the latter, indicating that for reconsolidation, the late 12-hour delay of sleep reconsolidated the reactivated old negative memory in the way of reducing false details, protecting them from the interference of updating. In a word, both true memory and false memory evolved across the delay with wakefulness and sleep after reactivation, but in distinct patterns, stating complicated reconsolidating mechanisms of different memory types. We infer that sleep plays an important role in memory reconsolidation, primarily via preserving less false memory. However, since we cannot wholly exclude the intermixed testing effect and do not set a wake group for control, the conclusions will need further exploration.

### Positive updating increases emotional arousal of aversive memories after 24-hour interval, including sleep

Apart from investigating memory alternation that results from emotional updating, we were also interested in altering the emotional significance of the negative memory during recall by inserting positive emotion when memory was reactivated and thus susceptive for modification. Our data showed, on Day 9, emotional arousal of the negative story had shown significant interaction effect, i.e., introducing neutral rather than positive emotional right after memory reactivation reduced the emotional arousal of the previous aversive story 24 hours later, while the valence of the aversive memory did not differ from each among four groups in our experimental manipulations. This result was seemingly counterintuitive at first sight. However, it could still be supported by previous literature. Although counter-conditioning has been shown effective to update addictive behaviors (Das et al., 2015; Goltseker et al., 2017), evidence suggested that after incomplete reminders, the omission of expected stimuli (i.e., extinction) could be more effective than introducing opposite valence. For example, in a study modifying alcohol-additional behavior in humans, prediction errors of omission (alcohol was unexpectedly removed) facilitated memory updating, rather than introducing the opposite feeling (alcohol was bitter than expected) (Hon, Das, & Kamboj, 2016). Furthermore, although studies show that introducing an opposite valence could reduce the initial negative conditioned response (e.g., pain movements (Claes, Karos, Meulders, Crombez, & Vlaeyen, 2014)), confounding results also exist. For instance, after reactivating the fear-conditioned animals, introducing counter-conditioning tended to prone even stronger fear response than an extinction procedure (Holmes et al., 2016). The idea of omitting emotional arousal benefits the distinction of aversive feeling has been utilized in clinical psychotherapies, for example, the *Systematic Desensitization* therapy, developed by Joseph Wolpe, in treating phobia (Wolpe, 1990). A typical protocol of this therapy involves a gradual exposure of the fearful stimulate. After each exposure, a relaxation process is introduced to help the patient prepare themselves for even stronger stimuli. The mechanism of this phenomenon was suggested that omitting expected aversive stimuli (in our case is the abruption of the negative story presentation) can lead to a rewarding result that is mediated by a prediction error signal in the mesocorticolimbic dopaminergic system (Kalisch, Gerlicher, & Duvarci, 2019; McNally, Johansen, & Blair, 2011). As a result, dopamine could accumulate in multiple brain regions, including the areas that are important for emotional memory like the amygdala (Bissière, Humeau, & Lüthi, 2003; Correia, McGrath, Lee, Graybiel, & Goosens, 2016; Marowsky, Yanagawa, Obata, & Vogt, 2005) and the hippocampus (Broussard et al., 2016; Chen et al., 2019; Ramirez et al., 2015). To note, at the time when prediction error was generated was also the time when the negative memory was reactivated. Hence, the relatively high dosage of dopamine in the brain could likely increase the emotional aspect of the reactivated memory (Luo et al., 2018). It was known that dopamine is also released during emotional processing (Badgaiyan, Fischman, & Alpert, 2009), therefore introducing positive emotion might maintain the high release of dopamine, thus possibly strengthening the emotional aspect of the negative memory. On the contrary, neutral updating materials would not trigger more dopamine, hence benefits a quick reduction of dopamine in memory reactivation brain regions. To note, we are not concluding that it was exactly the valence difference of the updating material that determined the emotional arousal on Day 9 morning because the arousal related neuromodulators (e.g., norepinephrine) also naturally fluctuate during sleep (Mitchell & Weinshenker, 2010), which might co-work on the modulation of the emotional arousal of the reactivated memory. However, since all experimental groups underwent a normal nocturnal sleep, we would thus speculate the valence of updating materials should at least modulated the emotional arousal of the negative story on Day 9.

Thus far, we are not aware of any studies showing the changes in subjective emotional arousal ratings after aversive-to-appetitive counter-conditioning. What we could compare was the studies with the measure of skin conductance response (SCR), since the subjective ratings of arousal using Self-Assessment Manikin (SAM) positively correlate with the magnitude of skin conductance response (Lang, Greenwald, Bradley, & Hamm, 1993). One fMRI study using monetary reward to counter-conditioning electrical shocks had failed to find statistically meaningful effects on SCR between a rewarded versus neutral updating material (Bulganin, Bach, & Wittmann, 2014). However, the current research was not entirely the same as that study in experimental paradigms. While the previous study followed the traditional Pavlovian conditioning procedure and gave the counter-conditioning phase not long after the fear acquisition, our study is established on emotional episodic memory, and it applied the positive intervention when the initial memories are completely consolidated after seven days. Thus, the different emotional updating might disturb the original emotional arousal differently while our study followed the reactivation procedure to make the consolidated memory labile change. Our study also provided evidence that the emotional valence rating, unlike the emotional arousal result, did not show any significant difference on Day 9. This was not surprising because emotional valence and arousal are supported by different brain regions. As it was shown in an fMRI study, the detection of valence was mainly supported by prefrontal cortex-hippocampal networks, whereas the emotional arousal mainly depends on amygdala-hippocampal networks (Elizabeth A. Kensinger & Corkin, 2004). Hence, it seems that valance evaluation reflects the semantic knowledge of emotion types, while arousal truly reflects emotional responses (Elizabeth A Kensinger & Corkin, 2003; Lang et al., 1993; Russell, 1980). Besides, it is understandable that one is unlikely to perceive it as either neutral or positive when facing a story about injury and illness because our knowledge of emotion types determined so.

## Conclusions

The current study investigated a behavioral method to modify aversive memory, benefiting from memory reconsolidation processes. We have provided novel evidence to disentangle the pros and cons of combining memory reactivation and emotional updating for different aspects of consolidated aversive memory. Our results suggest that the memory reactivation approach can significantly impair the true memory of the aversive story, which effect maintained over a 24-hour interval. On the contrary, false memory that represents the degree of memory distortion is augmented when the counter emotion (i.e., positive) was attached right after the old memory is reactivated only during the first 12-hour wake interval, but it attenuated after a night of sleep. Moreover, combining memory reactivation and emotional updating could also result in unwanted side effects, for example, hindering the reduction of emotional arousal for the aversive feelings. Our results provided important indications for the development of memory intervention strategies in real-world applications: the exact therapeutic methods should be carefully evaluated and tailored to each individual’s situation. For example, when reducing the aversive feelings of an already happened trauma is more urgent, the therapist could select the reactivation + neutral updating strategy, whereas when the subjective feeling of the old memory is less important than reconstructing the original memory to introduce more positive outcomes, the reactivation + positive updating strategy could be considered. However, for the latter case, how to prevent the weakening effect from sleep should be carefully investigated by future studies.

## Methods and Materials

### Participants

Eighty left, right-handed young adults (mean age ± s.e.m., 21 ± 0.255 years ranged from 18 to 27, 65 female subjects) with normal or corrected-to-normal vision participated in this study. Participants followed a regular daily work-rest schedule and reported no sleep-related problems during the last two months. All participants reported a good quality sleep during the night after the Day 8 Recall, with 7 – 9 h sleep (mean ± s.e.m 7.16 ± 0.078). Informed written consent was obtained from all participants before the experiment, and the Institutional Review Board approved the study protocol for Human Subjects at Beijing Normal University. Participants were assigned into four experimental groups, with each group of twenty subjects. We have controlled the gender bias, resulting in 13 males and 7 females in each experimental group. There was also no significant difference in age across all four groups.

All participants reported not to nap habitually nor have any sleep disorders (e.g., sleep apnea, irregular sleep, and insomnia) and did not take any medication at the experimental appointments. None of the participants claimed to have known neurological and psychiatric disorders (such as attention deficit hyperactivity disorder, anxiety, and depression). Participants had an age-appropriate sleep-wake rhythm (7-9 hours night/day), and they were not on any night shifts during the whole experimental procedure. Participants were instructed to keep a regular sleep schedule, abstain from caffeine- and alcohol-containing drinks on the experiments’ days.

### Experimental Procedure

The whole experiment took nine days in total, with a 7-day interval after the initiate encoding. On Day1, the encoding session started around 8 a.m.-10 a.m. Participants encoded one negative story consisted of 11 episodic slides with paired audio narratives, analog to previous studies (Cahill, Prins, Weber, & McGaugh, 1994; Kroes et al., 2014) (**Figure 1**). Participants were not instructed to remember anything. Instead, they were told to watch the slides carefully and meanwhile, pay attention to the auditory narratives. After encoding, participants took a 10-min rest. After the 10-min rest, participants were asked to recall the encoded negative story slide-by-slide at their own pace, with the instruction *Please describe the No. X slide as detailed as possible*. After this immediate recall session, participants left the lab and did not know what they would have to do when returning to the lab after one week.

We asked all the participants to come back to the lab again for delayed recall seven days later. We designed to update the old memory seven days later because previous studies showed that the longer time the memory develops, the more susceptible it is to new information (Scully, Napper, & Hupbach, 2017). Participants in the reactivation groups were instructed to recall the learned negative story slide-by-slide seven days before (see **Memory Reactivation**), while the other two Non-reactivation groups were told to watch a new story (see **Episodic Stories**). All participants were asked not to nap during the daytime and then came back to the lab 12 hours later (around 21:00 of the same day) for the Delayed Recall I. The delayed recalls were the same as the Immediate Recall session. Afterward, participants were allowed to go home. They were instructed to keep their natural sleep schedule, and they should return to the lab the next morning.

About 12 hours later, at about 9 a.m., participants returned to the lab for Delayed Recall Session II, which was also the same as the other two recalls. At the end of the experiment, participants were told to score the valence and arousal of the negative story again using the same procedure as the encoding.

### Episodic Stories

During the initial encoding session, participants were shown a negative story with 11 slides on the PC screen, accompanied by an auditory narrative forming an episode (See **Figure 1** as an illustration). Pictures of all slides were searched from the internet with free copyrights. All images were 27 cm high, 20 cm wide, with 1280 × 1024 Pixels. A pre-recorded male voice read the narrative of the negative story, which volume was controlled to 50 dB for the whole presentation. The negative story outline was about “I” observing a severe traffic accident, in which a young boy injured his leg seriously, and he has been sent to a hospital for a stressful operation and leaves severe sequelae eventually. The negative story was the same for all experimental groups.

During the updating session, participants learned a new story (either positive or neutral) identical in structure and scene to the negative story they saw in the encoding session. Of note, to guarantee that it is the emotion rather than the complication of the visual contents, the updating stories used the same images for visual cues, but we played different emotional narratives (positive or neutral) to describe the episodes of the presented slides **(Figure 8).** While the neutral story was in plain and ungarnished words describing neutral events on the pictures, the positive story talked about “me” vising a city and have experienced and observed a series of exciting and funny events **(Figure 8).** Both the positive and neutral narratives were from reading from a female voice, the volume of which is also controlled to 50 dB.

**Figure 7:**
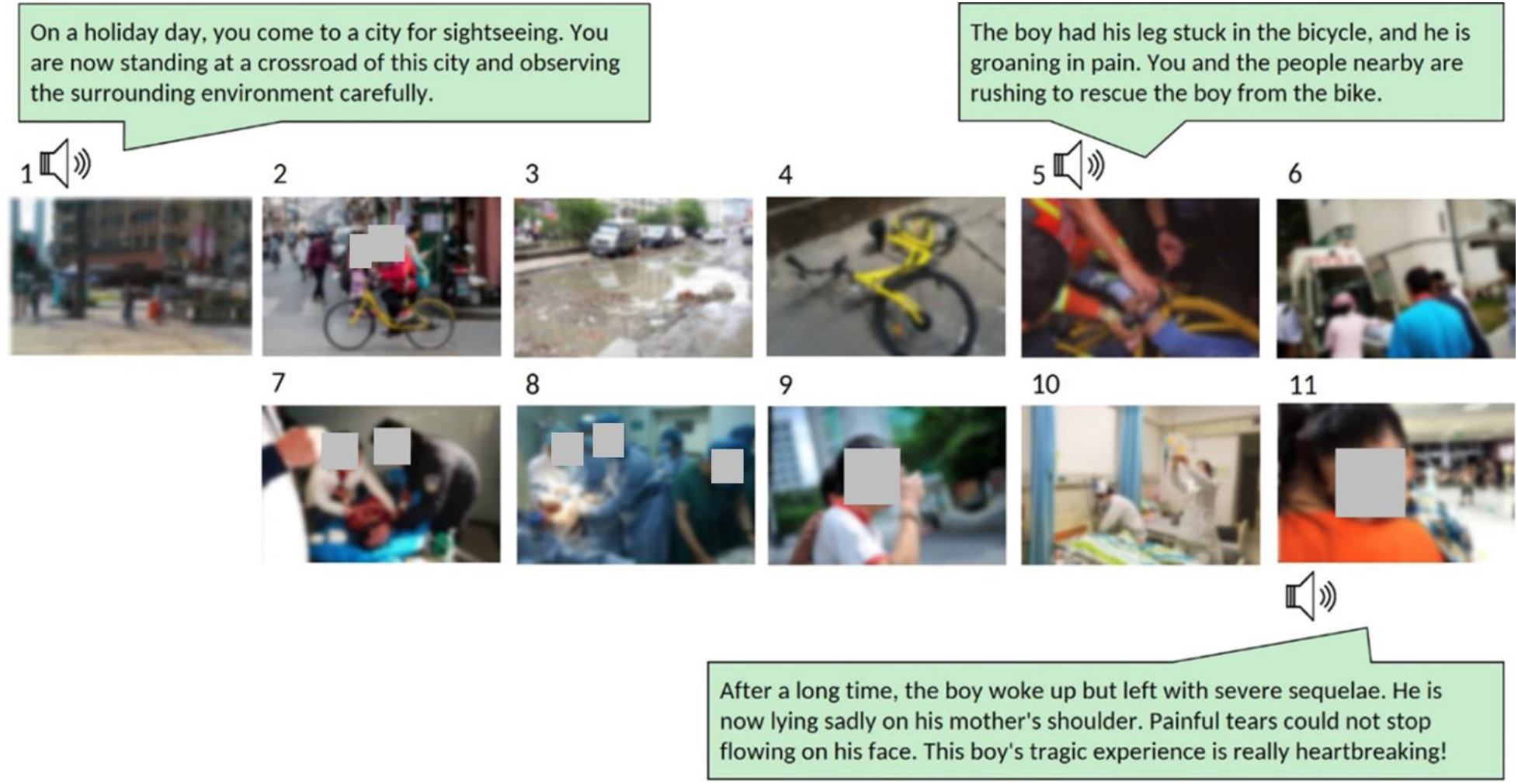
The Encoding Story. All participants were presented with an emotionally negative story consisting of 11 slides on PC, with auditory narratives of each slide from a middle-aged male voice. The whole story lasted about six minutes.

**Figure 8:**
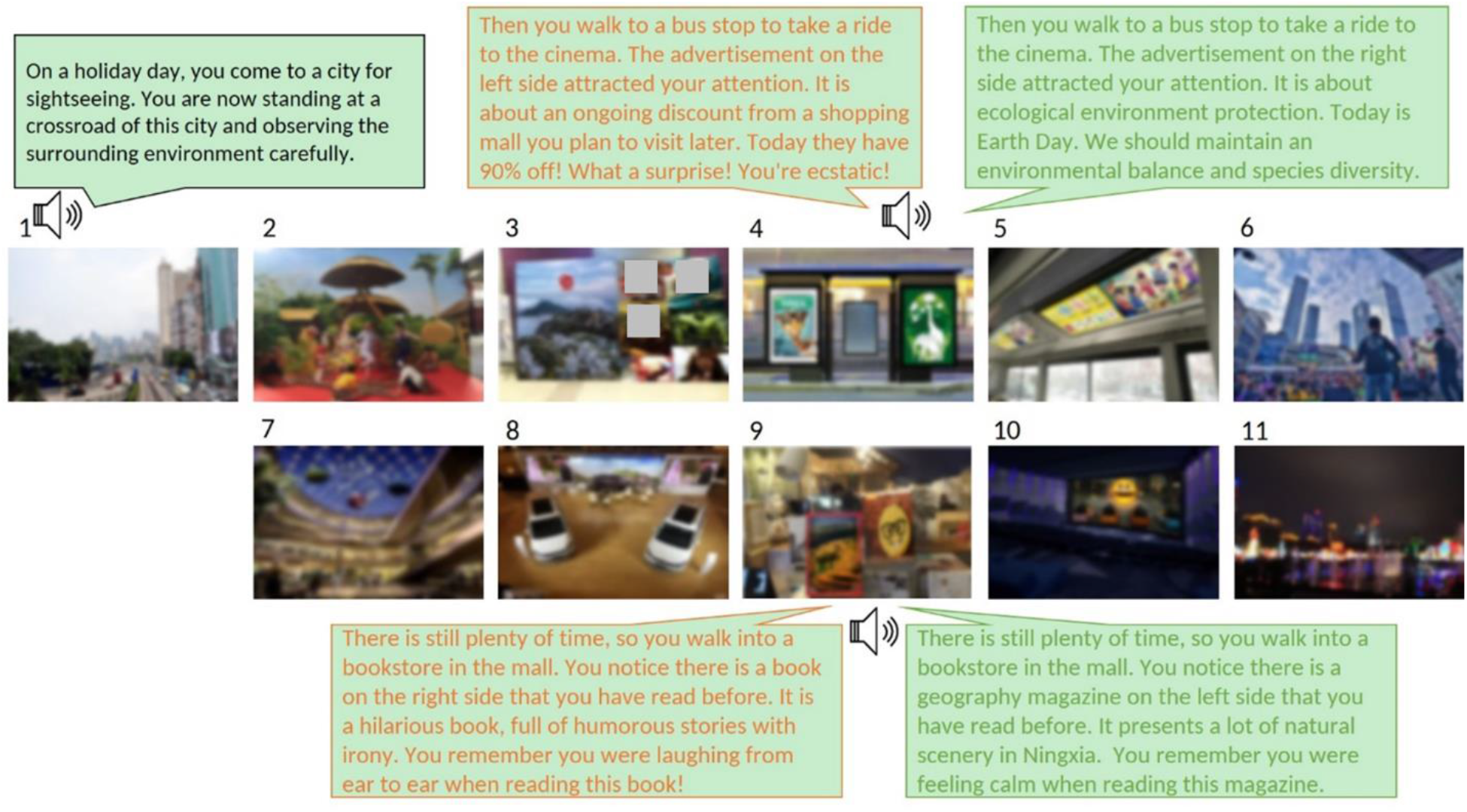
The Updating Stories. Participants in Positive and Neutral groups were presented with either a positive or a neutral story, respectively. All participants viewed the same pictures but with different auditory narratives (except the first slide). Both stories consisted of 11 slides, and each slide was accompanied by an auditory narrative from a young female voice. Orange: the positive narrative example, Green: the neutral narrative example. Each story lasts about six minutes.

We validated the emotional valence of three episodic stories used in this study. The subjective rating of the negative story’s valence and arousal for all participants were 2.190 ± 0.046 and 5.540 ± 0.178, respectively. There were no significant differences across four groups (all p > 0.75). The valence of the positive (3.790 ± 0.082) and neutral story (3.380 ± 0.052) was significantly different from each other (p < 0.001, Mann-Whitney’s test). The arousal of the positive and neutral stories was 4.655 ± 0.259 and 4.166 ± 0.259, respectively. However, the arousal of the two stories did not reach significant level (t(78) = −1.330, p = 0.187).

### Memory reactivation

Memory reactivation operations were adopted from previous studies that show successful memory labilizations (Forcato, Fernandez, & Pedreira, 2014). In particular, we introduced a prediction error that has been suggested to be critical for memory labilizations (Sinclair & Barense, 2019). Hence, for the participants of the reactivation groups, the reactivation session started by showing a partially masked first slide of the negative story. Participants were asked to answer three forced-choice questions related to the masked content. Once participants chose their answer, the corresponding mask was then removed to see the real picture as feedback. The reactivation score was calculated as the number of correctly answered questions by multiple choice. After the three questions, an unexpected black screen was suddenly presented, and the experimenter pretended to be surprised about the error and took the device away for solutions. The participants were instructed to relax for a few minutes without any distractions, like looking at their phones. Ten minutes later, the experimenter informed the participant with regret that the program was not fixable. Therefore the recall session had to be canceled, and the procedure continued to the next session, which was to watch a new story. The new story (either positive or neutral for different updating groups) consisted of another 11 slides was presented in the same way as the encoding of the negative story.

### Recall memory test

Three free recall tests were conducted 10 min after encoding of the first day, 12 hours after the updating – the evening of the eighth day, and the morning of the ninth day. Before the recall test, participants need to recall according to a suggested framework, including the location, characters, events in the story, and the shape, clothes, and actions of the characters. The participants did not receive any feedback during the whole process of free recall. To ensure all related memories are reported, if the participants ended their recall within 3 minutes, they are encouraged by the experimenter to describe more, such as “Please describe the detail in this slide as much as you can” and “Is there anything else you can recall?” If participants reported nothing more, the experimenter continued to ask the next slide. This procedure continued until all eleven slides were recalled.

Each day’s memory performance was first scored for each slide according to each participant’s oral report on correct (as True Memory) or incorrect information (as False Memory) of the negative episodic story. The final memory score was calculated as the sum of all slides, without the first slide that has been used for memory reactivation (i.e., the sum of slides 2 to 10). True memory was scored as any correct description of the corresponding slide. If the description was accurate (e.g., the boy was wearing a red jacket), this information was scored as 2 (i.e., one score for the “boy” and one score for the “red jacket”). If the description was not accurate but also not wrong (e.g., “there are many doctors perform surgery on children,” instead of “there are six doctors perform surgery on children”), this information was then scored as 0.5. False memory was scored according to any false information that the participant recalled about the corresponding image, including the incorrect details (e.g., wrong colors of someone’s cloth, wrong location of an object, and et al.), and created information that was not presented on the slides.

### Emotion self-assessment test

For each story presentation, i.e., the encoding of the negative story on Day1, the positive/neutral updating stories on Day8, and the final recall of the negative story on Day9, participants rated their subjective feeling of the slides on valence and arousal. The rating system was adopted from a standard emotion assessment instrument: The Self-Assessment Manikin and the Semantic Differential (SAM). Participants were told to understand their feelings as much as possible, and then to score their real valence and emotional arousal level, for 5 points scoring (1 = very negative, 2 = relatively negative, 3 = neutral, 4 = relatively positive, 5 = very positive) and 9 points scoring (the higher the score is, the stronger the arousal level), respectively. Emotion related score has been calculated as the absolute performance of Day 9, introducing their rates on Day1 as a covariate variable to control the pre-test effect.

### Statistical analyses

Unless it is stated otherwise, statistics for testing group differences for memory performance, subjective emotion rating, and control measures were all first submitted to 2 (Reactivation) × 2 (Updating) × 2 (sleep: before vs. after sleep) repeated ANOVA or 2 (Reactivation) × 2 (Updating) ANOVA. When a significant interaction effect or main effect was found, *post hoc* tests followed significant ANOVA effects, including Student’s *t*-test, if variances were unequal, or Welch’s *t*-test with an approximation for the degrees of freedom, or Mann–Whitney *U* test if normality of both samples was not violated. Both true memory and false memory were scored as the memory percentage on Day B relative to Day A (e.g., Day 8/ Day 1, Day 9/ Day 1, Day 9/ Day 8). Values of group descriptive statistics were presented as Means ± Standard Error, unless stated otherwise. Effect sizes for ANOVAs are partial eta squared, referred to as η^2^p. We used an alpha level of .05 for all statistical tests. Memory performance of all recalls is calculated based on ten slides, excluding the first slide.

Emotional valence and arousal were calculated as the mean value of all slides. To note, the validity of emotion for all three stories was the mean of all 11 slides, whereas the emotion ratings on both Day 1 and Day 9 were calculated as the mean of 10 slides (excluding the cue slide) to avoid any unexplainable influence from the slide that has been viewed extra in the reactivated groups. Two-group comparisons of the emotional valence and arousal were tested with parametrical independent samples *t*-test and nonparametric Mann–Whitney *U* test.

## Acknowledgements

This work was supported by the National Natural Science Foundation of China (31522028, 81571056), the Open Research Fund of the State Key Laboratory of Cognitive Neuroscience and Learning (CNLZD1503), and the Fundamental Research Funds for the Central Universities. We thank Ruiyi Chen and Liping Zhuang for their assistance in pilot experiments.

## Author contributions

S.Q., J.W. and C.F. conceived the experiment. B.C. performed data collection. B.C., J.W., and M.S. performed data analysis. J.W. drafted the manuscript, with B.C. & M.S. drafted the methods and results. All authors wrote the manuscript, contributed to data discussion and interpretation.

## Competing interests

The authors declare no competing interests.

## Notes

### Competing Interest Statement

The authors have declared no competing interest.

